# Learning and attention reveal a general relationship between neuronal variability and perception

**DOI:** 10.1101/137083

**Authors:** Amy M. Ni, Douglas A. Ruff, Joshua J. Alberts, Jen Symmonds, Marlene R. Cohen

## Abstract

The trial-to-trial response variability that is shared between pairs of neurons (termed spike count correlations^1^ or *r*_SC_) has been the subject of many recent studies largely because it might limit the amount of information that can be encoded by neuronal populations. Spike count correlations are flexible and change depending on task demands^2-7^. However, the relationship between correlated variability and information coding is a matter of current debate^2-14^. This debate has been difficult to resolve because testing the theoretical predictions would require simultaneous recordings from an experimentally unfeasible number of neurons. We hypothesized that if correlated variability limits population coding, then spike count correlations in visual cortex should a) covary with subjects’ performance on visually guided tasks and b) lie along the dimensions in neuronal population space that contain information that is used to guide behavior. We focused on two processes that are known to improve visual performance: visual attention, which allows observers to focus on important parts of a visual scene^15-17^, and perceptual learning, which slowly improves observers’ ability to discriminate specific, well-practiced stimuli^18-20^. Both attention and learning improve performance on visually guided tasks, but the two processes operate on very different timescales and are typically studied using different perceptual tasks. Here, by manipulating attention and learning in the same task, subjects, trials, and neuronal populations, we show that there is a single, robust relationship between correlated variability in populations of visual neurons and performance on a change-detection task. We also propose an explanation for the mystery of how correlated variability might affect performance: it is oriented along the dimensions of population space used by the animal to make perceptual decisions. Our results suggest that attention and learning affect the same aspects of the neuronal population activity in visual cortex, which may be responsible for learning- and attention-related improvements in behavioral performance. More generally, our study provides a framework for leveraging the activity of simultaneously recorded populations of neurons, cognitive factors, and perceptual decisions to understand the neuronal underpinnings of behavior.

---

We investigated the relationship between the activity of neuronal populations and behavioral performance by manipulating attention and perceptual learning simultaneously in two rhesus monkeys (*Macaca mulatta*). We recorded from neuronal populations in visual area V4 with chronically implanted microelectrode arrays as the monkeys practiced an orientation change-detection task that manipulated spatial attention (**Fig. 1a**). The monkey’s task was to detect a subtle change in the orientation of either of two Gabor stimuli that flashed on and off simultaneously, one at a location that overlapped the receptive fields (RFs) of the recorded neurons and the other in the opposite hemifield (**Fig. 1b**).

Before recording, the monkeys were briefly trained on the behavioral task so that they were familiar with the task structure. Each monkey was trained to report 90° changes with the attention cue in place: the cued stimulus would change on 80% of trials and the uncued stimulus would change on 20% of trials. We began recording after 2-5 days of training on the full version of the task (see Methods), with performance on cued trials matched between the two stimuli.

We designed our experiment to allow us to simultaneously measure attention and perceptual learning in the same behavioral trials and neuronal responses. We quantified the behavioral effects of attention (comparing attended trials (RF stimulus cued) and unattended trials (RF stimulus uncued) within each session) or learning (across sessions) by quantifying the monkey’s detection sensitivity for a fixed orientation change (*d’*; other behavioral measures gave qualitatively similar results; see **Supplementary Fig. 1**) at the RF location (**Fig. 1c**).

Attention and perceptual learning affected performance in similar ways. Both processes were associated with improvements in behavioral sensitivity (**Fig. 2a**, **h**), albeit on different timescales (attention is the difference between the solid and dashed lines in **Fig. 2a** and **h**, and learning is the change in performance across time). Consistent with other perceptual learning paradigms^21,22^, the behavioral improvements associated with learning were specific to the trained stimulus location (**Supplementary Fig. 2**).

**Figure 1.**
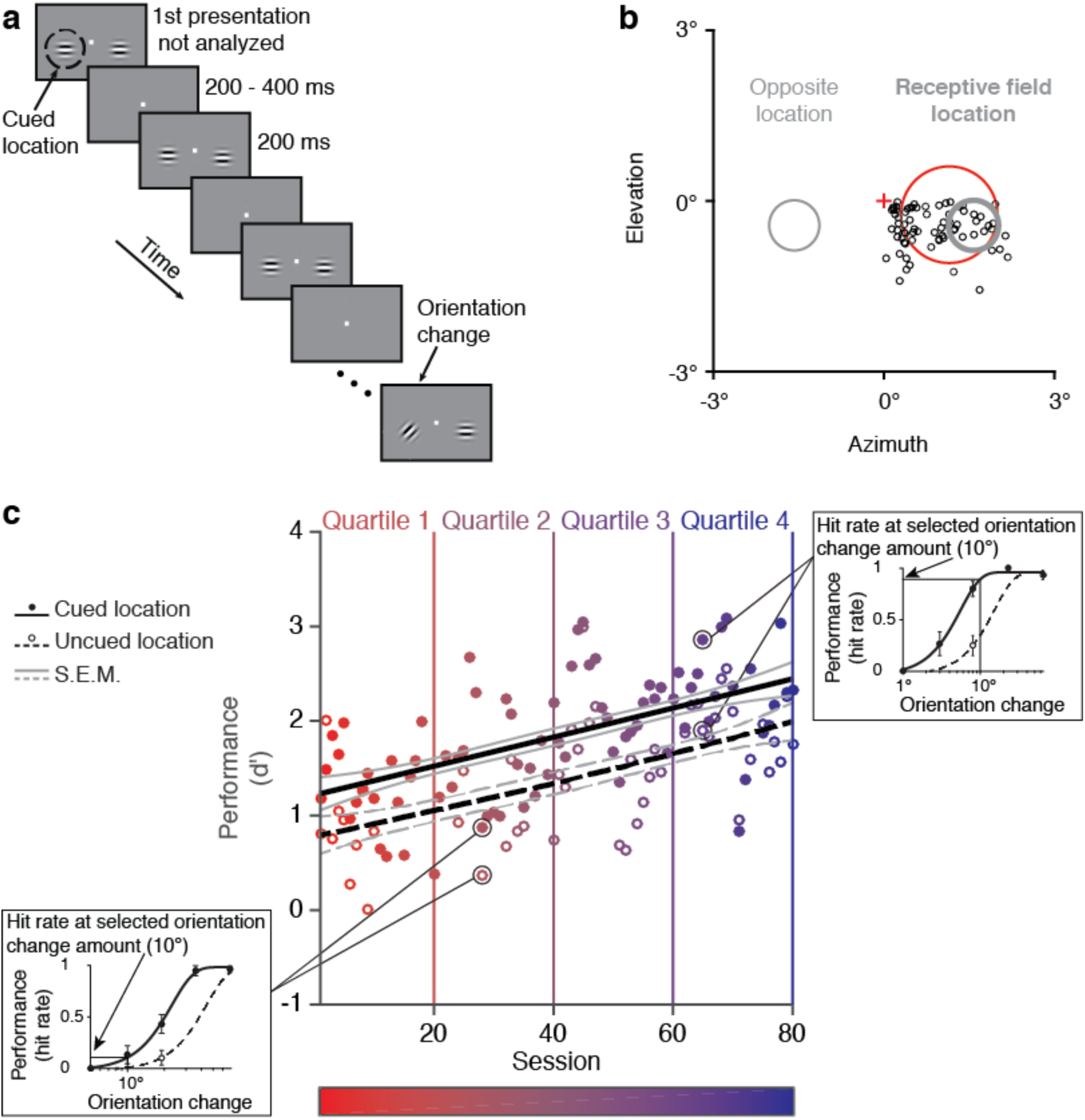
Methods and behavior. **a**, Orientation change-detection task with cued attention^2^. **b**, Centers of visual receptive fields for the recorded units from one monkey (*black circles*). The monkey fixated a central point (*red cross*) while two Gabor stimuli were presented, one overlapping the neuronal receptive fields (*thick gray circle*) and one in the opposite hemifield (*thin gray circle*). The red circle illustrates a representative receptive field size. **c**, Our method for quantifying attention- and learning-related changes in detection sensitivity (*d’*) as a function of session number (one session = 125 trials in each attention condition; multiple sessions per day; see Methods). The best fitting exponential functions are plotted for cued (*solid black line* fit to *filled circles*) vs. uncued (*dashed black line* fit to *empty circles*) performance, with S.E.M. indicated (cued: *solid gray lines*; uncued: *dashed gray lines*). The heat map illustrates the session number and learning quartiles, which we used throughout the paper to illustrate learning phase. Insets: Psychometric curves (hit rate as a function of orientation change amount) for two example sessions to illustrate how we calculated hit rate at one selected orientation change amount for each animal (Monkey 1: 29°, Monkey 2: 10°).

Both attention and perceptual learning had profound effects on the variability of neuronal responses in V4. Even though the trial-averaged evoked response of individual units did not change consistently with learning^21,23-27^ (**Fig. 2b**, **i**), attention and learning were associated with decreases in the mean-normalized trial-to-trial variance (Fano factor) of individual units and the correlated variability (*r*_SC_) between pairs of units (**Fig. 2c**, **d**, **j**, **k**; see **Supplementary Fig. 3** for eye movement analysis). These decreases in variability appear to be task-specific, as variability in the responses of the same neurons to visual stimuli presented during passive fixation was constant throughout the recording period (**Fig. 2f**, **g**, **m**, **n**). The data also suggest that the monkeys had already learned to attend during the initial training period and they were improving sensitivity at orientation change-detection rather than learning to attend during the recording period. The perceptual learning-related increase in behavioral sensitivity across sessions was not accompanied by changes in the signatures of attention across sessions (**Supplementary Fig. 4**).

Task-related changes in response variability (particularly correlated response variability) have attracted considerable attention because they have the potential to change the amount of sensory information encoded in a population of neurons, which might limit performance. In theoretical studies, however, the relationship between response variability and population coding is a matter of active study^8-11,28,29^. The fundamental concern is that correlated variability should only affect information coding if it lies along the dimensions in neuronal population space along which task related information is read out^10,14^. However, determining whether the correlated variability would affect an optimal decoder based on the thousands of neurons that respond to any stimulus is nearly impossible with experimentally tractable data sets.

**Figure 2.**
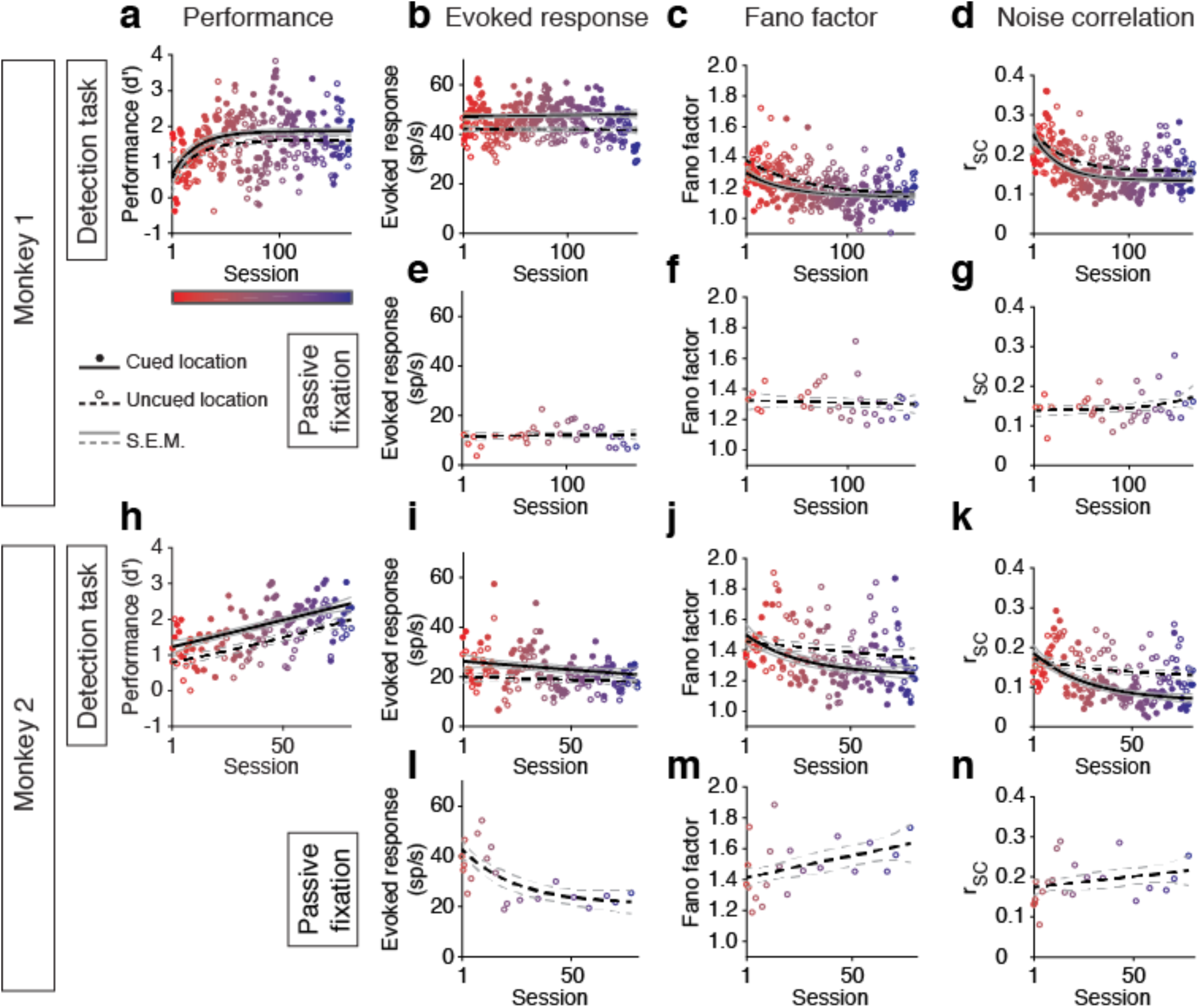
Summary of the behavioral and neuronal effects of attention and perceptual learning. Each plot follows the format of **Fig. 1c**. We quantified the effects of attention using a paired t-test comparing cued and uncued trials within each session and the effects of learning during the cued attention condition using a two-tailed t-test comparing sessions from the first vs. second half of the total training period, without assuming equal variances. Number of sessions: detection task: Monkey 1: *n* = 150, Monkey 2: *n* = 78; passive fixation: Monkey 1: *n* = 35, Monkey 2: *n* = 22. Mean number of single or multiunits per session: Monkey 1: *n* = 34, Monkey 2: *n* = 15. **a-g**, Monkey 1. **h-n**, Monkey 2. **a,h**, Sensitivity (*d’*) increased with both attention (Monkey 1: *p* < 10^-^ 8; Monkey 2: *p* < 10^-12^) and learning (Monkey 1: *p* < 10^-4^; Monkey 2: *p* < 10^-9^). **b,e,i,l**, Evoked response (firing – baseline rate) increased with attention (Monkey 1: *p* < 10^-37^; Monkey 2: *p* < 10^-^14) but did not change consistently with learning or passive fixation (no change in Monkey 1, *p* = 0.13 learning, *p* = 0.44 fixation; decrease in Monkey 2, *p* < 10^-3^ learning, *p* < 10^-3^ fixation). **c,j**, Fano factor decreased with both attention (Monkey 1: *p* < 10^-5^; Monkey 2: *p* < 10^-4^) and learning (Monkey 1: *p* < 10^-5^; Monkey 2: *p* < 10^-3^). **f,m**, Fano factor decreased only in the context of the detection task and not during passive fixation (Monkey 1: *p* = 0.14; Monkey 2: *p* = 0.05). **d,k**, Correlated variability decreased with both attention (Monkey 1: *p* < 10^-8^; Monkey 2: *p* < 10^-8^) and learning (Monkey 1: *p* < 10^-4^; Monkey 2: *p* < 10^-5^). **g,n**, Correlated variability did not change during passive fixation (Monkey 1: *p* = 0.47; Monkey 2: *p* = 0.47).

We addressed the importance of attention- and perceptual learning-related changes in response variability by investigating their relationship to behavior. One strong prediction of the hypothesis that response variability limits task performance is that changes in response variability should always be associated with changes in psychophysical performance, regardless of whether the changes in variability came about from attention, learning, or some other factor.

Consistent with this prediction, we found that there is a single, robust relationship between correlated variability and perceptual performance, whether changes in perceptual performance happen quickly (attention) or slowly (learning; **Fig. 3a,b**). This relationship does not simply reflect the long-term changes in correlated variability and performance due to perceptual learning or the changes caused on a faster timescale by attention. It also reflects factors outside experimental control: the relationship between correlated variability and detection sensitivity was robust even when we examined the residuals of each measure after removing the variability captured by the exponential fits in **Fig. 2** (**Fig. 3c,d**). These results show that correlated variability in visual cortex is a reliable indicator of performance in this task.

We used two additional, complementary measures of population activity to further investigate the hypothesis that the attention- and perceptual learning-related decreases in response variability were responsible for the behavioral improvements we observed. First, we calculated the ability of an optimal, cross-validated linear decoder to detect changes in the orientation of the stimuli we presented. In small neuronal populations, decreases in correlated variability would be expected to reduce redundancy and increase the information encoded by the population if the neurons responded similarly to the stimulus change^6,30^. In fact, in our data set, the vast majority of the units fired more strongly in response to the orientation change (93% of units; presumably because of a release from adaptation).

**Figure 3.**
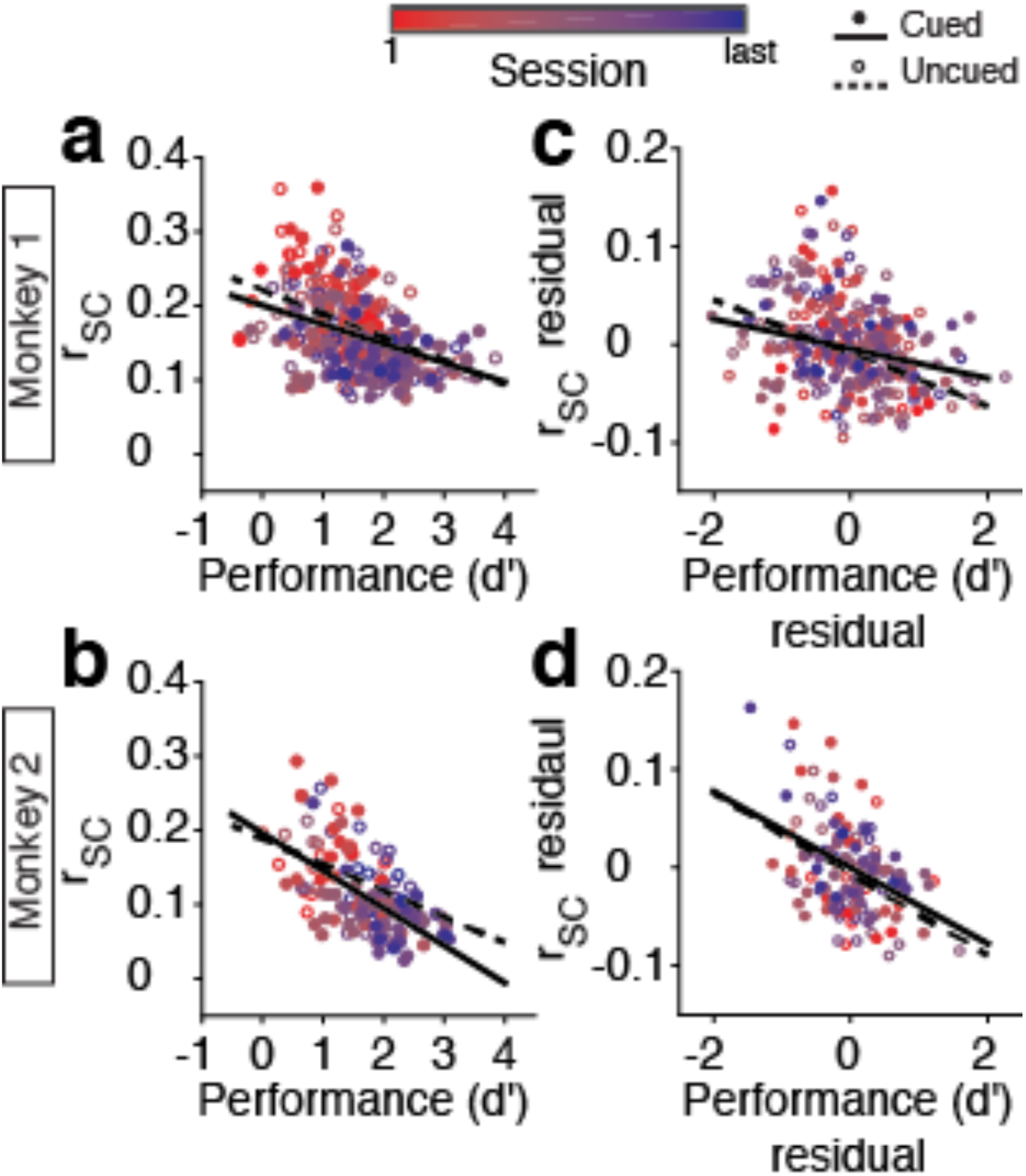
The relationship between correlated variability and performance is the same for attention and perceptual learning. **a,b**, Mean *r*_SC_ and *d’* were significantly correlated for both animals and the relationship between the two was indistinguishable for the two attention conditions. **a**, Monkey 1: Pearson correlation coefficients: cued: *R* = −0.40, *p* < 10^−7^; uncued: *R* = −0.48, *p* < 10^−9^; the cued and uncued correlation coefficients were indistinguishable (ZPF test: *zpf* = −0.92, *p* = 0.36). **b**, Same for Monkey 2. Cued: *R* = −0.59, *p* < 10^−9^; uncued: *R* = −0.45, *p* < 10^−4^; *zpf* = 1.12, *p* = 0.26. In both animals, the relationship between Fano factor and *d’* was weaker (data not illustrated). Monkey 1 cued: *R* = −0.15, *p* = 0.08; uncued: *R* = −0.30, *p* < 10^−4^. Monkey 2 cued: *R* = −0.56, *p* < 10^−7^; uncued: *R* = −0.28, *p* = 0.06. **c,d**, The relationship between *r*_SC_ and *d’* persisted even when we looked only at the residuals after removing the effects of attention and learning (exponential fits in **Fig. 2**). **c**, Monkey 1: *r*_SC_ residual vs. *d’* residual. Cued: *R* = −0.26, *p* < 10^−3^; uncued: *R* = −0.40, *p* < 10^−7^; *zpf* = 1.5, *p* = 0.12. **d**, Same for Monkey 2. Cued: *R* = −0.45, *p* < 10^−5^; uncued: *R* = −0.44, *p* < 10^−3^; *zpf* = 0.04, *p* = 0.97. Number of sessions: Monkey 1: *n* = 150, Monkey 2: *n* = 78. To test whether the residuals contained attention- or learning-related trends not captured by the exponential fits, we ran an ANOVA per monkey to test the effects of session number and attention condition on the *d’* residual, and an ANOVA per monkey to test the effects of those same two variables on the *r*_SC_ residual: we did not find any significant main effects or interactions for either monkey (*p* > 0.40).

Both attention and perceptual learning improved the performance of the optimal stimulus decoder (**Fig. 4a-d**). The attention- and learning-related differences in decoder performance tended to increase with increasing number of units (the lines in **Fig. 4a-d** diverge), suggesting that changes in the relationships between multiple units, rather than changes in the means or variability of the responses of single neurons, were responsible for the improvement in decoder performance. Consistent with this idea, the relationship between correlated variability and the amount of information encoded by the neuronal population was the same for attention and learning (**Fig. 4e-f**).

Although it is tempting to infer from the results in **Fig. 4** that attention and perceptual learning improve the amount of visual information encoded in V4, the neuronal populations we recorded are small subsets of the neurons that encode task-relevant visual information, and it is possible that changes in correlated variability do not affect the amount of visual information encoded in larger populations. However, the robust relationship between correlated variability, the amount of information encoded by small populations, and behavior suggests that correlated variability is at least a byproduct of the process causally responsible for improving performance.

We reasoned that we could examine the relationship between correlated variability and performance more directly by looking at the relationship between population activity and the animal’s behavior on a trial-by-trial basis. For example, finding that correlated variability can predict errors would imply a close relationship between spike count correlations and decisions. However, comparing variability to individual choices requires a measure of correlated variability on a single trial, and spike count correlations (and Fano factor) are only defined over many trials. We therefore used principal component analysis (PCA) on population responses to repeated presentations of the same visual stimulus (the stimuli before the orientation change; the same stimuli used to compute spike count correlations in **Fig. 2-3**) to identify the axis in population space that accounts for the most correlated variability. To do so, we plotted the responses of the population in a population space where each dimension represents the firing rate of one unit, and performed PCA on the result (**Fig. 5a**). Because this cluster of points consisted of population responses to repeated presentations of the same visual stimulus, the first PC represents the dimension that accounts for the most shared trial-to-trial variability across the population (dashed line in **Fig. 5a**). Consistent with the recent observation that correlated variability is typically low dimensional^31-34^, we found that the variance explained by the first PC accounted for the majority of the session-to-session variability in spike count correlations, even when we accounted for the changes caused by attention and perceptual learning (**Fig. 5b,c** and **Supplementary Fig. 5**).

**Figure 4.**
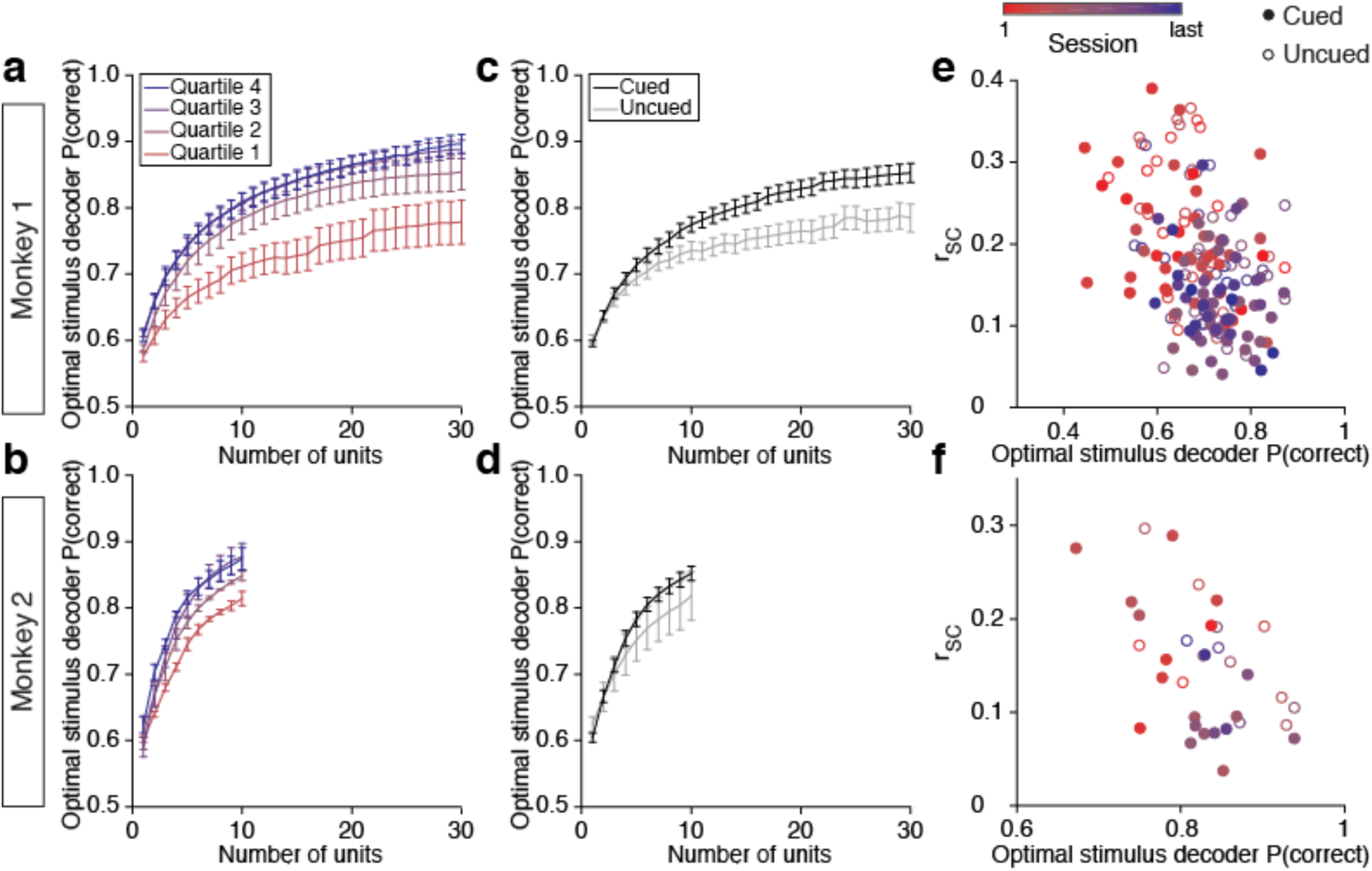
The ability of an optimal, linear, cross-validated decoder to detect changes in the visual stimulus improves with perceptual learning and attention in a way that is predicted by changes in correlated variability. **a**, Optimal stimulus decoder performance improves throughout learning over a long time period (see **Fig. 1c** for learning quartile illustration) for Monkey 1, **b**, and Monkey 2, **c**, as well as with attention within each day for Monkey 1, **d**, and Monkey 2. Error bars are S.E.M. Number of days: Monkey 1: *n* = 37, Monkey 2: *n* = 10. **e**, The relationship between correlated variability (*r*_SC_) and optimal stimulus decoder performance is the same for attention and learning for Monkey 1: Pearson correlation coefficients: cued: *R* = −0.41, *p* < 10^−5^; uncued: *R* = −0.37, *p* < 10^−3^; ZPF test: *zpf* = 0.30, *p* = 0.77, **f**, and Monkey 2: cued: *R* = −0.56, *p* = 0.01; uncued: *R* = −0.66, *p* = 0.01; *zpf* = −0.44, *p* = 0.66. The relationship between Fano factor and decoder performance was weaker (Monkey 1 cued: *R* = −0.19, *p* = 0.06; uncued: *R* = −0.18, *p* = 0.15. Monkey 2 cued: *R* = −0.53, *p* = 0.02; uncued: *R* = −0.42, *p* = 0.14). Number of sessions/days (days separated into sessions when possible): Monkey 1: *n* = 101, Monkey 2: *n* = 20.

These analyses show that, to a first approximation, variability along the first PC accounts for pairwise spike count correlations. This puts us in a position to assess the importance of correlated variability to the monkey by determining whether population activity along this first PC can predict the monkey’s choices on a trial-by-trial basis.

We found that activity along this first PC (and therefore correlated variability) has a much stronger relationship with the monkey’s behavior than its influence on the performance of the stimulus decoder. A linear, cross-validated choice decoder (**Fig. 5a**) could detect differences in hit- vs. miss-trial responses to the changed stimulus as well from population activity along the first PC alone as it could maximally differentiate hit- vs. miss-trial responses with larger numbers of PCs (green lines; **Fig. 5d,e**). In contrast, the stimulus decoder (**Fig. 5a**) was much worse at detecting differences in responses to the previous stimulus (the stimulus prior to the change) vs. the changed stimulus based on the first PC alone as compared to its maximum performance with larger numbers of PCs (black lines; **Fig. 5d,e**).

**Figure 5.**
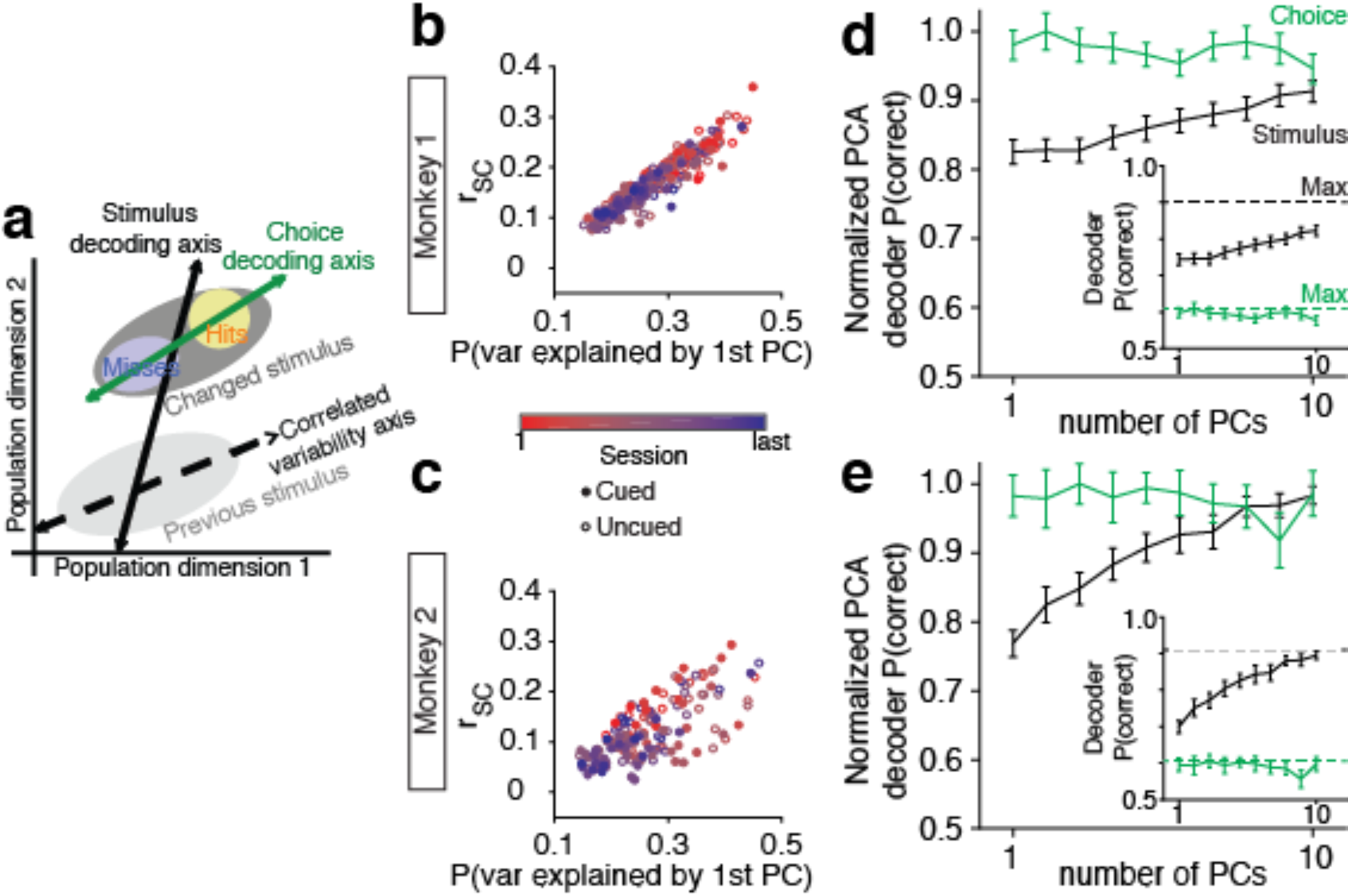
Correlated variability affects the monkey’s behavior more than it affects the optimal stimulus decoder. **a**, Decoder schematic. Responses to the stimulus before the change (previous stimulus, *light gray ellipse*) and to the changed stimulus (*dark ellipse*) are plotted in a subset of a high dimensional space in which each dimension represents the responses of one unit. The axis that explains the most variability in responses to the previous stimulus (*dashed line*) is by definition the axis that explains the most correlated variability. The choice decoder (*green line*) decodes differences in responses between detected (hits) and missed stimulus changes (*yellow and blue ellipses*). The stimulus decoder detects differences between the neuronal responses to the previous and changed stimuli. **b,** The mean *r*_SC_ is highly correlated with the proportion of variance explained by the first PC (*dashed line* in **a**). Monkey 1: Pearson correlation coefficients: cued: *R* = 0.94, *p* < 10-68; uncued: *R* = 0.94, *p* < 10-68. The relationship remains for the residuals of the exponential fits for *r*SC vs. variance explained by PC 1. Cued: *R* = 0.92, *p* < 10^−60^; uncued: *R* = 0.92, *p* < 10^−61^. **c**, Same for Monkey 2. Cued: *R* = 0.67, *p* < 10-11; uncued: *R* = 0.67, *p* < 10-11. Residuals: cued: *R* = 0.62, *p* < 10-9; uncued: *R* = 0.67, *p* < 10-11. Number of sessions: Monkey 1: *n* = 150, Monkey 2: *n* = 78. **d**, PCA choice decoder and PCA stimulus decoder performance per number of PCs, normalized to the respective decoder’s maximum performance (each decoder was run with all testable numbers of PCs, from 1 to a maximum of 42). Inset shows raw decoder performance. For Monkey 1, the PCA choice decoder could distinguish hit from miss trials as well from population activity along the first PC only (*leftmost point of the green line*) as it could maximally distinguish with larger numbers of PCs (*p* = 0.44). The stimulus decoder could not distinguish the previous from the changed stimuli as well based on the first PC (*leftmost point of the black line*) as it could maximally distinguish with larger numbers of PCs (*p* < 10^−3^). Error bars are S.E.M. **e**, Same format as **d**, for Monkey 2. Choice decoder paired t-test: *p* = 0.93. Stimulus decoder paired t-test: *p* < 10-6. Number of days: Monkey 1: *n* = 37, Monkey 2: *n* = 10.

Unsurprisingly for a population of neurons in visual cortex, a linear decoder could detect the stimulus change from the neuronal population responses (black lines; **Fig. 5d,e** insets) much better than it could detect the animal’s choices from those same responses (green lines; **Fig. 5d,e** insets). However, the relative influence of the first PC was much stronger for the choice decoder than for the stimulus decoder. We normalized the performance of each PCA decoder per number of PCs to its own maximum performance to highlight the very different slopes (**Fig. 5d,e**). The choice-predictive activity was essentially completely explained by variability along the first PC, while the stimulus-predictive signals along the first PC were much lower than their peak. The choice decoder uses the monkey’s choices to infer the most important subspace of population activity for the monkey’s decisions, and this subspace was highly influenced by correlated variability.

It is difficult to determine from extracellular recording data whether choice-predictive signals come from a bottom-up, causal relationship between sensory responses and decisions or from trial-to-trial variability from cognitive factors or post-decision signals^35,36^, and a recent study identifying the directionality of choice-predictive signals in mouse sensory cortex found that they are both bottom-up and top-down in origin^37^.

To determine whether the choice-predictive activity in the populations of neurons we recorded is well positioned to causally affect decisions, we examined the time course of the choice-predictive activity. Neuronal responses to the changed stimulus were calculated based on each neuron’s initial response to the changed stimulus (60-130 ms after stimulus onset, which corresponds to the first 70 ms of the evoked response after the response latency of V4 neurons) to avoid artifacts from behavioral responses (the monkeys began eye movement responses to the changed stimulus an average of 210 ms after stimulus onset; as a note, all changed and previous stimulus responses were taken from this same time frame for all decoder analyses). We compared the choice-predictive activity in the first half of this time frame (60-95 ms) to that of the second half (96-130 ms) and found that the choice-predictive activity was as strong during the first spikes of the stimulus response (Monkey 1: mean of 61% correct decoder performance; Monkey 2: mean of 60%) as it was later in the response (Monkey 1: mean of 60%, paired t-test: *p* = 0.43; Monkey 2: mean of 57%, *p* = 0.25). That the choice-predictive activity described here is present early in the evoked response suggests that it does not reflect post-decision feedback. Therefore, while we cannot determine whether the choice-predictive signals come from sensory or cognitive factors, they are present during the full decision-making period.

The results from **Fig. 5** are consistent with the idea that correlated variability influenced the monkeys’ performance. This would mean that the monkeys are suboptimal in a very particular way, such that correlated variability strongly influences performance. To investigate whether the monkeys’ choices were influenced by activity along the first PC (and thus, spike count correlations; **Fig. 5b,c**) in a complementary way, we compared projections of population responses to the stimuli before the orientation change onto the first PC with weighted sums of population activity using a method described by Haefner and colleagues^9^ to infer the weights the monkeys used to make decisions (based on the correlation structure and the neuronal responses to the changed stimulus on hit vs. miss trials). We found that the projections onto the first PC were correlated with the weighted sums predicted by this decoding method for both monkeys (Monkey 1: median Pearson correlation coefficient across days: *R* = 0.69; two-tailed Wilcoxon signed rank test of the Pearson correlation coefficient across days: *p* < 10^−8^; Monkey 2: *R* = 0.48, *p* < 10^−6^). Together, these results suggest that while an optimal decoder may in theory be able to discount correlated variability, the monkey’s choices can be predicted by the very dimension that is most influenced by correlated variability.

## Discussion

We showed that attention and perceptual learning have the same effects on populations of neurons in visual cortex, and that changes in spike count correlations might underlie behavioral improvements. Correlated variability covaries with performance in ways that are indistinguishable for attention, learning, and factors outside experimental control (based on comparisons of residuals that exclude attention and learning effects in **Fig. 3c,d**), and population activity along the dimension that explains most correlated variability is strongly associated with the animal’s choices on a trial-by-trial basis.

The notable perceptual learning-related changes in spike count correlations we observed are in contrast to the often modest effects of learning on the activity of single neurons that we and others observed. Most prior electrophysiological studies of perceptual learning that focused on the trial-averaged activity of single neurons found, as did we, minimal to no effects of learning on evoked firing rates in visual cortex^21,23-27^. A study of pairs of simultaneously recorded units found that spike count correlations varied across subjects based on training experience, but did not find a relationship between this shared variability and population coding efficiency^5^, while other studies suggest that learning shapes neuronal population measures^38,39^. These results are consistent with the idea that correlated variability might affect decision-making through means other than the information that can be gleaned by an optimal decoder. Our approach allowed us to study perceptual learning in two ways: measuring its effects on neuronal populations and comparing it to visual attention in the same neurons and trials. This approach revealed that attention and learning have similar effects on visual cortex, including indistinguishable effects on spike count correlations that are well positioned to affect performance.

An alternative hypothesis is that the monkeys were learning to attend throughout the recording period, and that the behavioral and neurophysiological effects of attention and perceptual learning were similar because perceptual learning acts through attention^18-20,23,25,26,40^. However, the effects of attention did not change throughout the perceptual learning period, as the behavioral and neuronal signatures of attention did not change across sessions (**Supplementary Fig. 4**).

The robustness of the relationship between correlated variability and perceptual performance, whether detection sensitivity changes on a moment-by-moment basis due to shifts in attention or gradually over long periods through perceptual learning, suggests that while the mechanisms of attention and learning act on different time frames, these processes share a common computation in terms of their effects on the information encoded in visual cortex. Some characteristics of this computation are informed by recent studies showing that a low rank modulator whose strength is affected by attention could simultaneously account for the attention-related changes in rate, Fano factor, and correlated variability in populations of V4 neurons^33-34^. An intriguing possibility proposed by a recent theoretical study^32^ is that attention and perceptual learning decrease the strength of such a modulator by changing the balance of inhibition and excitation in V4. Such a mechanism might improve performance through some combination of improving the amount of visual information encoded in populations of neurons and improving the fidelity with which that information is communicated to the downstream areas involved in forming perceptual decisions^12,41^. Studying how very different processes such as attention and learning affect perception in common ways provides a new framework for understanding the relationship between neuronal population activity and perception.

Spike count correlations have been a subject of many studies in part because they provide a tempting explanation for why performance on sensory tasks is worse than the amount of information encoded by neuronal populations with independent neurons^14^. Spike count correlations are flexible and change depending on the behavioral task in ways that seem consistent with the hypothesis that they limit performance on psychophysical tasks^2-7^. However, the relationship between correlated variability and population coding is complicated because it depends strongly on population size, and determining whether spike count correlations could change the information encoded by large populations would require simultaneous recordings from an experimentally unfeasible number of neurons over an even more impossible number of behavioral trials^10,14^.

We approached this question using behavior as our anchor, and found two lines of evidence suggesting that spike count correlations affect psychophysical performance. First, there is a robust, consistent relationship between correlated variability and performance, which is identical for attention and perceptual learning. Second, correlated variability is associated with the animals’ choices on a trial-by-trial basis: variability along the axis in population space that was most closely associated with spike count correlations accounted for essentially all of the choice-predictive activity in our recorded population of neurons.

These results suggest that 1) if the change in correlated variability does not cause the improvements in performance associated with attention and perceptual learning, it is a byproduct of the neuronal mechanism that does and 2) the decision-making mechanism is suboptimal in a way that emphasizes the impact of correlated variability. This might arise because some biological mechanism (perhaps related to the aspects of population activity that are communicated to the downstream neurons involved in perceptual decision making^41^) causes correlated variability to affect the readout of neuronal populations more strongly than predicted by an optimal decoder. It is difficult to dissociate whether the monkeys are acting optimally with less information or suboptimally: in the future, inactivation experiments may help make this distinction^11,42^. Our results suggest that correlated variability is well posed to limit performance on visually guided tasks.

## Methods

The subjects were two adult male rhesus monkeys (*Macaca mulatta*, 8 and 10 kg). All animal procedures were approved by the Institutional Animal Care and Use Committees of the University of Pittsburgh and Carnegie Mellon University.

### Behavioral task

Before recording, we trained each monkey on the basic orientation change-detection task (**Fig. 1a**) and the meaning of the attention cue. Attention was cued to the left stimulus in one block of 125 trials, and to the right stimulus in a second block, with the two blocks making up a single session. In each session, 20% of trials (all at the middle or largest orientation change) were invalidly cued^2^. We began recording once the monkey’s behavior was stable enough to produce reliable fits of the Weibull function to the psychometric data. The size, location, and spatial frequency of the Gabor stimuli were fixed throughout learning. The orientation of all stimuli before the orientation change was consistent throughout each recording session but changed by 15° between days.

### Recordings

We recorded extracellularly from single units and sorted multiunit clusters (the term “unit” refers to either; 19-42 units per session, mean 34 for Monkey 1; 6-25 units per session, mean 15 for Monkey 2) in V4 of the left hemisphere using 96-channel microelectrode arrays (Blackrock Microsystems) as previously described^2^. We presented visual stimuli and tracked eye position as previously described^6^.

The data presented are from 42 d of recording for Monkey 1 and 28 d of recording for Monkey 2. Each day consisted of 1-7 sessions (mean of 3.6/d for Monkey 1; 2.9/d for Monkey 2), for a total of 150 sessions for Monkey 1 and 78 sessions for Monkey 2. Data were collected during passive fixation on 35 d for Monkey 1 and 22 d for Monkey 2.

### Data analysis

We based most neuronal analyses on spike count responses between 60-260 ms after stimulus onset. All analyses used correct and miss trials only (i.e., trials in which an orientation change occurred). To minimize the impact of adaptation on our results, we did not analyze the first stimulus presentation in each trial.

We only analyzed a recorded unit if its stimulus-driven firing rate was significantly higher than baseline (Wilcoxon signed rank test; *p* < 10^−10^). We only included complete sessions, and excluded sessions from analyses if average baseline activity across included units was less than 20 Hz, and outlier sessions were excluded from analyses based on the Tukey method.

We fit sets of data across all sessions with the following exponential equations. For exponential decay of increasing form:

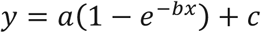

For exponential decay of decreasing form:

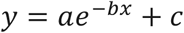

We compared the correlation between two variables in the cued attention condition to the correlation between the same two variables in the uncued attention condition using the ZPF test for dependent but non-overlapping Pearson’s correlation coefficients^43^.

### Decoder

The optimal stimulus decoder was a linear classifier with leave-one-out cross validation that was trained to discriminate the stimulus before the change from the changed stimulus. We measured decoder performance as a function of population size. The maximum number of units per monkey was based on classifier constraints on the pooled covariance matrix (Monkey 1: 30 units; Monkey 2: 10 units). We randomly selected subsets of units without replacement 1000 times for each population size. To maximize the number of behavioral trials, we analyzed all trials in a given day together (with the exception of the comparisons to spike count correlations and Fano factor in **Fig. 4e,f**, for which days were divided into sessions when possible), focusing only on trials that presented the middle orientation change amount, for which we had cued and uncued changes. Because the middle orientation change amount varied across recording days, we matched the distributions of orientation change amounts across learning in all analyses. For comparisons to spike count correlations and Fano factor (**Fig. 4e,f**), spike count correlations and Fano factor were calculated for the same stimuli used for the decoder.

To avoid artifacts in neuronal firing rates due to eye movements in response to the changed stimulus, we performed decoder analysis on the changed and previous stimulus responses with an abbreviated time window: spike count stimulus responses were measured between 60-130 ms after stimulus onset.

The PCA stimulus decoder differed from the optimal stimulus decoder only in that we decoded activity in the first *n* PCs instead of in the responses of subsets of *n* neurons. The PCs were based on responses to the stimulus before the orientation change as described in the text. All neuronal responses used for the decoder (responses to the stimulus before the change and responses to the changed stimulus) were projected onto those PCs. The PCA choice decoder (*Choice decoding axis*; **Fig. 5a**) classified population responses to the changed stimulus on hit vs. miss trials projected onto the PCs from the stimulus before the change.

## Acknowledgements

We thank Karen McCracken for technical assistance and John H. R. Maunsell and Adam Kohn for comments on a previous version of this manuscript. The authors are supported by US National Institutes of Health grants 4R00EY020844-03 and R01 EY022930 (M.R.C.), a Whitehall Foundation grant (M.R.C.), a Klingenstein-Simons Fellowship (M.R.C.), a Sloan Research Fellowship (M.R.C.), a McKnight Scholar Award (M.R.C.), and a fellowship (A.M.N.) and a grant from the Simons Foundation (M.R.C.).

## Author Contributions

A.M.N., D.A.R., and M.R.C. designed the experiments, A.M.N., D.A.R., J.J.A., and J.S. collected the data, A.M.N. performed the analyses, and A.M.N. and M.R.C. wrote the paper.

## Author Information

The authors declare no competing financial interests. Correspondence and requests for materials should be addressed to M.R.C.

